# SETD2 regulates the methylation of translation elongation factor eEF1A1 in clear cell renal cell carcinoma

**DOI:** 10.1101/2020.10.26.354902

**Authors:** Robert Hapke, Lindsay Venton, Kristie Lindsay Rose, Quanhu Sheng, Anupama Reddy, Angela Jones, W. Kimryn Rathmell, Scott Haake

## Abstract

SET domain-containing protein 2 (*SETD2*) is commonly mutated in renal cell carcinoma. SETD2 methylates histone H3 as well as a growing list of non-histone proteins. To explore SETD2-dependent regulation of the kidney cancer proteome, we performed a systems-wide analysis of protein lysine-methylation and expression in wild type (WT) and *SETD2-*knock out (KO) kidney cells. We observed decreased lysine methylation of the translation elongation factor eEF1A1. *EEF1AKMT2* and *EEF1AKMT3* are known to methylate eEF1A1, and we show here that their expression is dependent on SET-domain function of *SETD2*. Globally, we observe differential expression of hundreds of proteins in WT versus *SETD2-*KO cells, including increased expression of many involved in protein translation. Finally, we observe decreased progression free survival and loss of EEF1AKMT2 gene expression in *SETD2-*mutated tumors. Overall, these data suggest that *SETD2-*mutated ccRCC, via loss of enzymetic function of the SET domain, displays dysregulation of protein translation as a potentially important component of the transformed phenotype.

## Introduction

Renal cell carcinoma (RCC) is among the top ten most common cancers in the United States, with more than 70,000 new patients expected to be diagnosed in the next year^1^. Within RCC, the most common histological subtype is clear cell renal cell carcinoma (ccRCC), occurring in approximately 75% of cases^2^. ccRCC tumors follow a shared pathway of molecular pathogenesis, beginning with loss of the short arm of chromosome 3 (3p)^3^. Within 3p are numerous tumor suppressor genes (TSGs) that are commonly deleted in ccRCC, including *VHL, PBRM1, SETD2*, and *BAP1*. These 3p TSGs are thought to follow the two-hit hypothesis, with subsequent mutation or hypermethylation resulting in loss of function of the second allele. This is most commonly seen with *VHL* (∼90% of cases^4^). Though the loss of *VHL* is considered to be necessary for disease progression, it is not sufficient^5^. This loss is usually followed by the loss of one or more chromatin modifiers, including *PBRM1, BAP1, SETD2*, or *KDM5C*^6^. In ccRCC, inactivating mutation of the histone methyltransferase *SETD2* occurs in roughly 12% of primary tumors, and is likely lost in >60% of metastatic tumors^7^, suggesting its loss of function is important for disease progression.

*SETD2* regulates chromatin structure by contributing to the histone code^8^. “Writers”, “readers”, and “erasers” constitute the histone code and contribute to the regulation of gene expression^9^. This code is primarily made up of various post-translational modifications (PTMs) applied to the histone tail and serve distinct purposes^8^. Some PTMs, such as the acetylation of histone 4 lysine (H4K12), are repressive marks, indicating a silenced gene. Other marks, such as the trimethylation of H3K36, are activation marks, indicating an actively transcribed gene. SETD2 is the only known methyltransferase capable of H3K36 trimethylation^10,11^ and influences diverse processes around transcribed genes including mRNA splicing^12^, cryptic transcription^13^, DNA repair^14^, chromosomal segregation^15^, and others. Despite its role in these numerous pathways, a full understanding of the mechanism whereby loss of SETD2 promotes tumor progression remains unclear.

Although protein methylation has predominantly been studied in the context of the histone code, the vast majority of the methylproteome exists on non-histone proteins^16^. Other processes regulated by methylation include RNA metabolism, cell cycle progression, apoptosis, and protein translation^16^. Recently, lysine methylation has been shown to regulate protein translation in cancer, with METTL13-mediated lysine methylation of eEF1A1 being required for KRAS-driven tumors^17^. Other work has determined that methylation is intimately involved with peptide synthesis proteins, including both ribosomal proteins and translation factors^18^. A key elongation factor in protein translation, eEF1A1, has 6 known lysine methylation sites^19^. However, how these sites regulate protein translation remains to be fully elucidated.

Many methyltransferases are known to have multiple targets^16^. Recently, SETD2 has similarly been shown to methylate multiple proteins, including alpha-tubulin^20^, Signal transducer and activator of transcription 1 (STAT1)^21^, and Enhancer of zeste homolog 2 (EZH2)^22^. Thus, we sought to perform a systems-wide analysis of *SETD2-*dependent changes to the tumor proteome. Here, we quantitatively measured changes in protein lysine-methylation and expression in wild type (WT) and *SETD2-*knock out (KO) human kidney cells using mass spectrometry. Through this, we identified a novel role for SETD2 in regulating the methylation of eukaryotic elongation factor 1A (eEF1A1) as well as increased expression of proteins required for protein translation. In patient tumors, we similarly observed up-regulation of protein translation genes in *SETD2-* mutated tumors. These data suggest a new mechanism whereby loss of *SETD2* influences cell function during the progression of cancer.

## Results

### Loss of *SETD2* results in decreased eEF1A1 methylation

In order to study SETD2-dependent changes to lysine methylation in kidney cells, we performed immunoprecipitation (IP) followed by liquid chromatography-tandem mass spectrometry (LC-MS/MS) to catalog methyl-lysine peptides in human SV40 immortalized proximal tubule kidney cells (HKC) either expressing endogenous protein or lacking SETD2 from a model generated by TALEN deletion (Fig. 1). The *SETD2*-knock out (KO) cell lines were previously created in our lab and are described elsewhere^23^. SETD2 status in HKC cell lines was validated by western blot (Supplemental Fig. 1). Protein extract without IP was analyzed for total protein expression. Peptide quantification was facilitated with the use of Stable Isotope Labeling with Amino acids in Cell culture (SILAC)^24^ with global analysis demonstrating a normal distribution of heavy (labeled) and light (unlabeled) proteins (Supplementary Fig. 2).

**Figure 1.**
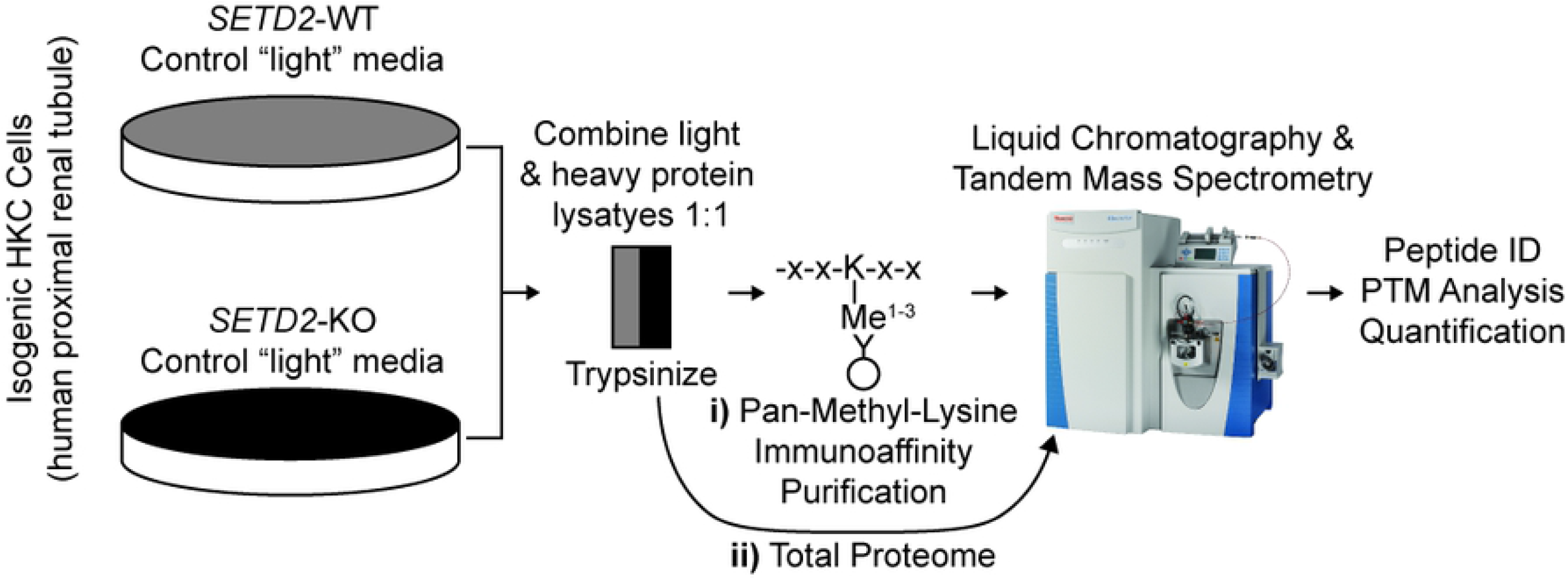
Workflow for the identification and quantification of peptides in wild type and *SETD2-*knock out human kidney cell lines. Above is the workflow for **i)** lysine-methylated (lysine-methylome) and **ii)** all peptides (total proteome).

**Figure 2.**
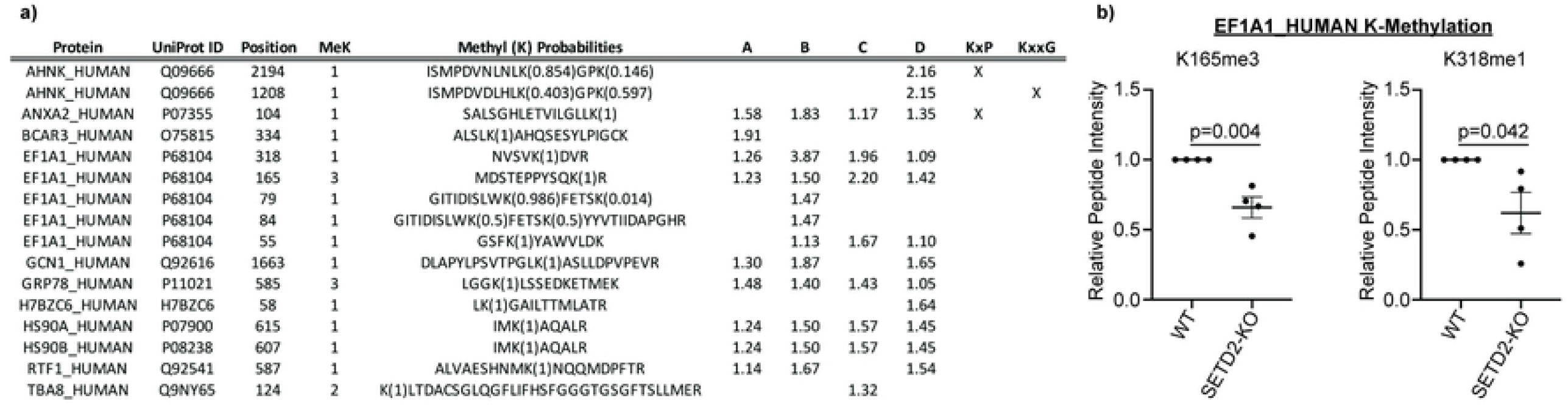
Peptides demonstrating loss of lysine methylation in *SETD2-*KO human kidney cell lines. **a)** “Position” corresponds to location of methylated lysine residue within the full-length protein. “MeK” indicates whether the lysine residue was mono-, di-, or tri-methylated. When more than one lysine was methylated within a peptide, “Methyl K Probabilities” indicates the relative probability that the indicated lysine residue was modified. A-D indicates the four biological mass spectrometry replicates and the ratio of the wild type (WT) and *SETD2-*knock out (KO) ratios (i.e., WT/KO ratio). If the lysine predicted to be modified corresponds to a canonical *SETD2* methylation motif (e.g., KxP or KxxG), it was indicated with “x”. Only peptides whose WT/KO ratio were >1 in all replicates and whose average was >1.25 are listed. See Supplementary Table 1 for the complete results. **b)** Relative intensities of the K165me3- and K318me1-containing peptides of the eEF1A1 protein in WT and *SETD2-KO* cells. Two-sided students t-test was used in this analysis.

Four biological replicates with SILAC-label swapping were performed for both the lysine-methylome and total proteome workflows, quantifying roughly 45 and 5000 proteins in each replicate, respectively (Table 1). Methylated lysine peptides whose WT/KO ratio was >1 (signifying decreased methylation in KO) in all measured replicates and whose average was >1.25 are listed (Fig. 2a, see Supplemental Table 1 for complete list of identified peptides). Multiple lysines in the eukaryotic translation elongation factor 1A1 (eEF1A1) were identified by these criteria. Specifically, trimethylation of eEF1A1 lysine(K)165 and monomethylation of K318 was significantly decreased in association with the loss of SETD2 protein (Fig. 2b).

**Table 1.** Total and lysine methylated peptides identified in wild type and *SETD2-*mutated human kidney cell lines.

| Replicate | A | B | C | D |
| --- | --- | --- | --- | --- |
| WT | H | L | L | H |
| KO | L | H | H | L |
| 2D/Total Protein |  |  |  |  |
| Total Proteins Quantified | 4863 | 5190 | 5002 | 5283 |
| Total Peptides Quantified | 27069 | 31773 | 29817 | 29800 |
| 1D/meK IP |  |  |  |  |
| Kme 1-3 Peptides Quantified |  |  |  |  |
| me1k | 37 | 33 | 29 | 39 |
| me2k | 14 | 14 | 11 | 9 |
| me3k | 12 | 10 | 11 | 10 |
| Kme 1-3 Proteins Quantified |  |  |  |  |
| me1k | 29 | 27 | 24 | 31 |
| me2k | 8 | 10 | 8 | 6 |
| me3k | 9 | 7 | 8 | 8 |

To validate that eEF1A1 lysine methylation was decreased in *SETD2-*KO cell lines, we evaluated the extracted ion chromatograms from the mass spectrometry data. This evaluation confirms that both K165 and K318 have decreased methylation in the *SETD2-*KO lysates (Fig. 3a & 3b). Next, we enriched eEF1A1 through a gel extraction of eEF1A1’s predicted migration distance (i.e., 50 kDa) from protein lysate that was not immunoprecipitated for lysine-methylated peptides. The K165 lysine methylated peptide abundance was compared in the SILAC-labeled protein lysates, again demonstrating decreased eEF1A1 K165me3 peptide in *SETD2-*KO cells (Fig. 3c). To confirm that decreased eEF1A1 methylation was not an artifact of SILAC labeling, we spiked gel extractions from non-SILAC labeled lysate with a synthetic heavy peptide containing the predicted K165 trimethylation. Using relative quantification, we again observe decreased K165 lysine methylated peptide with loss of SETD2 in HKC lysate (Fig. 3d). Finally, to rule out clonal effects, isogenic WT and *SETD2*-KO lysate from the human ccRCC cell line 786-O was tested using the previous workflow, similarly showing decreased K165me3 in *SETD2*-KO (Fig. 3d). Thus, in multiple cell lines and through multiple methods, we observe decreased eEF1A1 K165 and K318 methylation in *SETD2-*KO cells.

**Figure 3.**
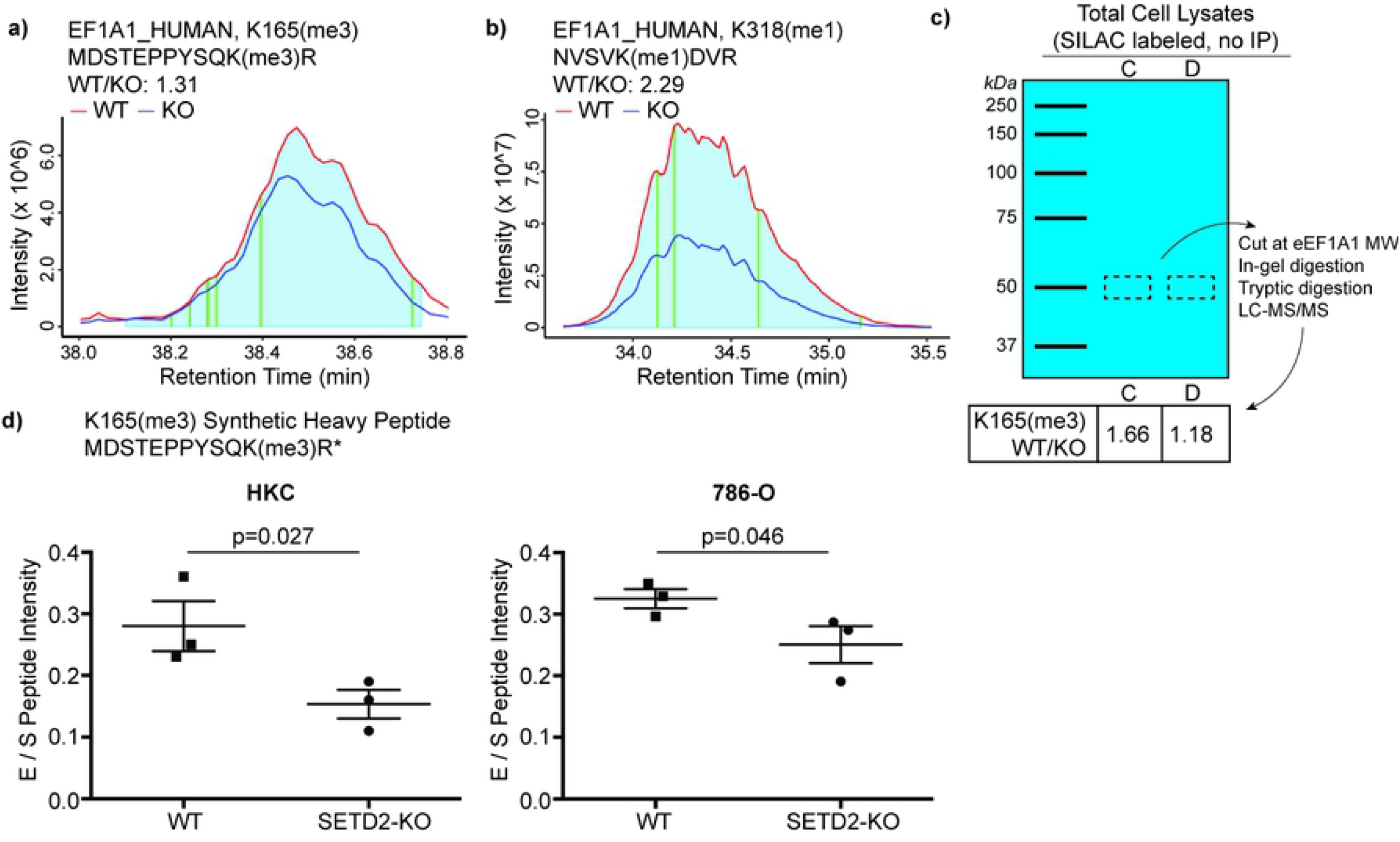
Loss of SETD2 expression results in loss of eEF1A1 methylation. In order to validate MaxQuant peptide quantification, the extracted ion chromatograms (EIC) were manually analyzed. The ratio of the **a)** eEF1A1_K165 peptide intensity was calculated in the WT and *SETD2-*KO lysates (AUC of the grey zones compared; vertical gold line corresponds to retention time at which the peptide was originally identified). **b)** EICs were also analyzed to validate peptide eEF1A1_K318 WT/KO ratio. **c)** The ratio of eEF1A1_K165 was compared in SILAC-labelled WT and *SETD2-*KO again. However, instead of enrichment via immunoprecipitation with anti-Kme(1-3) antibody, the eEF1A1 protein was cut from a gel (i.e., enrichment based on protein size) **d)** Synthetic peptide labeled with heavy arginine (13C, 15N) was formulated for eEF1A1_K165(me3). Synthetic peptides were spiked into WT and *SETD2-* KO lysates for HKC and 786-O, allowing for relative quantification of peptide in non-SILAC-labeled systems (one-sided t-test).

### Decreased eEF1A1 methyltransferase expression in *SETD2*-KO

While we have robust data that methylation of K165 and K318 on eEF1A1 decreases in *SETD2-* KO cells, neither lysine matches a known consensus sequence for SETD2-mediated lysine-methylation (i.e., KxP or KxxG)^10,20,21^. Furthermore, lysine methyltransferases responsible for these PTMs have previously been identified: EEF1AKMT3 methylates K165 and EEF1AKMT2 methylates K318^25–27^. Therefore, we sought to evaluate whether the expression of these enzymes was dependent on SETD2 protein. We observe that loss of SETD2 expression was associated with decreased expression of EEF1AKTM2/3 in HKC and 786-O cells as well as HEK-293T cells (human embryonic kidney cells) (Fig. 4a, b). To better understand the mechanism behind SETD2-dependent eEF1AKMT2/3 modulation, we rescued *SETD2*-KO cells with various mutated SETD2 constructs (Fig. 4c). These were designed by Hacker et al. and include a truncated but functional *SETD2* gene (WT-Tr), a SET-domain mutant (SET-mt), and an SRI-domain mutant (SRI-mt)^23^. The WT-Tr and SRI-mt constructs recover EEF1AKMT2/3 expression to wildtype levels, whereas the SET-mt does not (Fig. 4d, e). These data suggest that SET domain function of SETD2 is necessary for the expression of EEF1AKTM2/3 in human kidney and kidney cancer cells.

**Figure 4.**
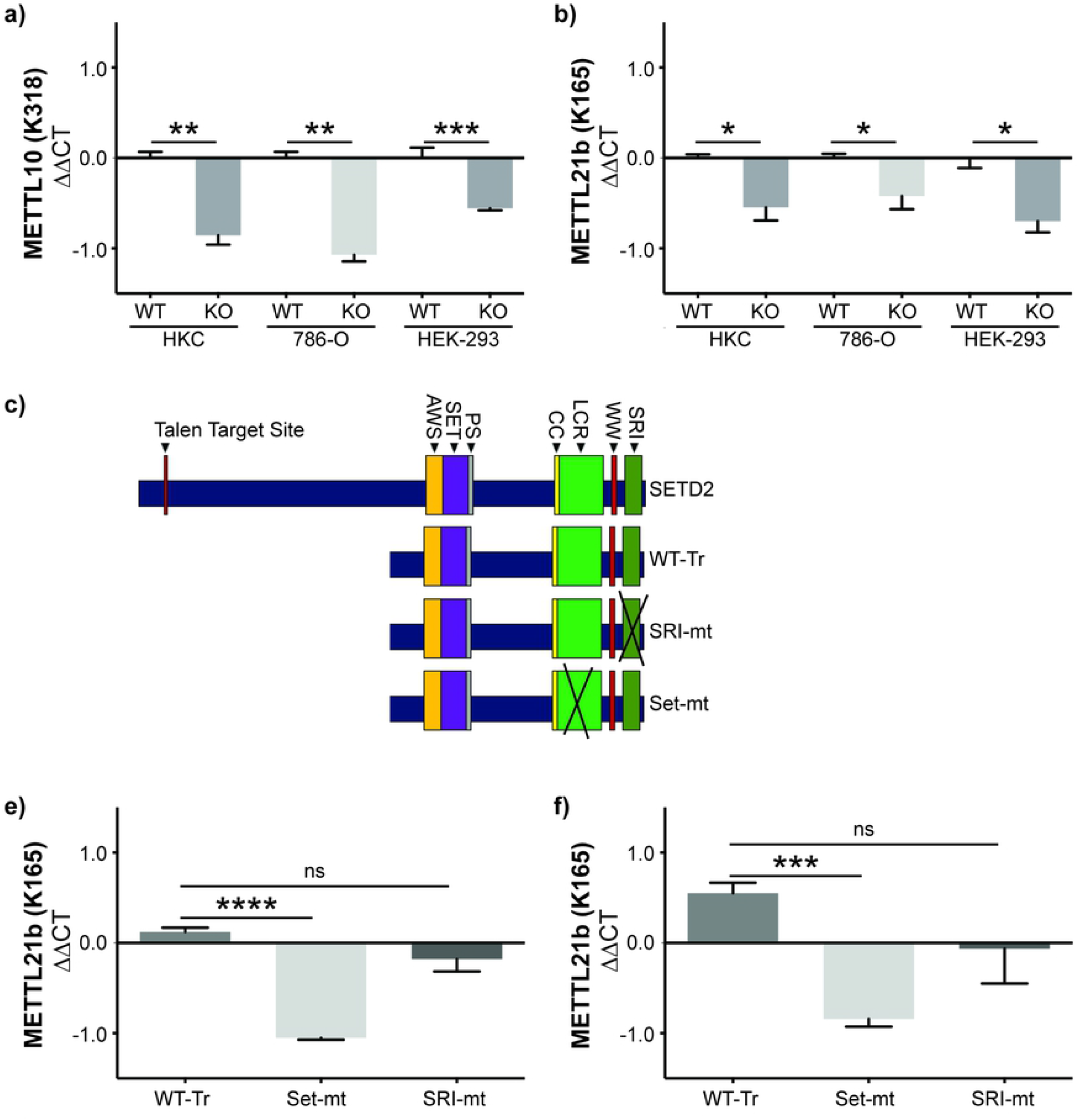
SET domain function of SETD2 is necessary for expression of methyl transferases that target eEF1A1_K165 and eEF1A1_K318. **a-b)** The expression of EEF1A1KMT2-3 was evaluated in WT and *SETD2-*KO human proximal tubule (HKC), clear cell renal cell carcinoma (786-O) and human embryonic kidney (HEK 293FT) cell lines **c)** The *SETD2-*KO HKC cell line was rescued with either a truncated WT *SETD2* (WT-Tr) or a truncated *SETD2* harboring mutations in the SRI (SRI-mt) or SET (Set-mt) domains (modified from Hacker et al., J Biol Chem 2016). **d-e)** The rescue cell lines were evaluated for gene expression of EEF1AKMT2 and EEF1AKMT3 via RT-PCR and normalized to *SETD2*-WT HKC cells. ns = p>0.05; *p<0.05; **p<0.01; ***p<0.001.

### System-wide changes to translation with loss of *SETD2*

Returning to the total proteome data, we observed significant system-wide changes in protein expression for wild type versus *SETD2-*KO HKC cells (Fig. 5a, Supplemental Table 2). WT/KO protein ratios were ranked and evaluated for Gene Set Enrichment Analysis (GSEA) using the Gene Ontology (GO) datasets. The most significant pathways to be increased in the SETD2-KO cell lines included GO_Translational_Initiation along with several other pathways pertaining to translation and/or ribosome function (Fig. 5b). Thus, along with decreased methylation of the translation elongation factor eEF1A1, our data suggests that loss of SETD2 results in increased levels of proteins associated with protein translation. Gene sets that are significantly decreased in the SETD2-KO cells include metabolic pathways, such as GO_NADH_Dehydrogenase_Complex (Fig. 5b). Shown is the enrichment plot for the most significantly increased pathway, GO_Translational_Initiation (Fig. 5c).

**Figure 5.**
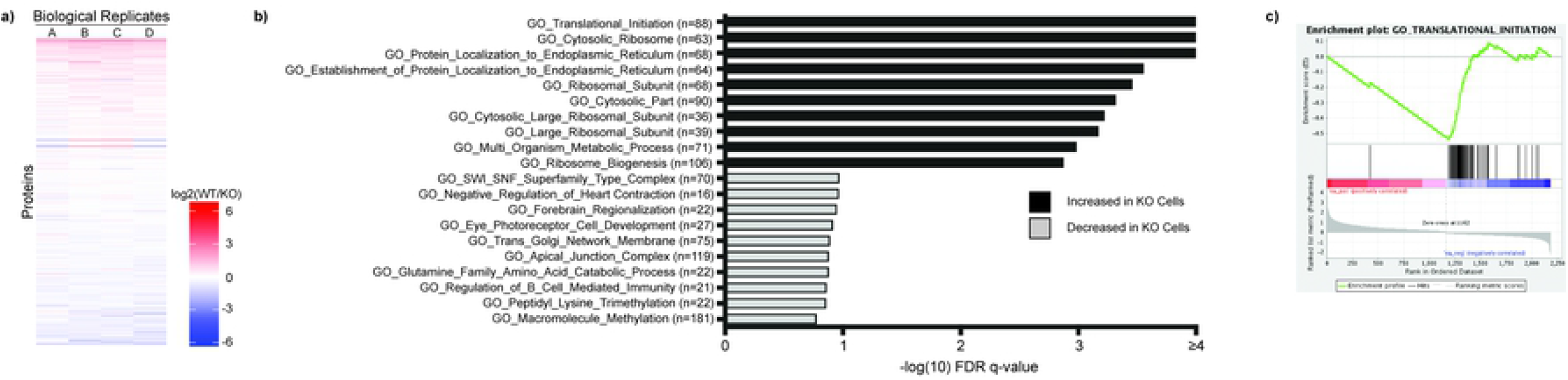
Total Proteome Analysis Demonstrates Changes in Protein Translation in *SETD2-*KO Cells. **a)** Individual proteins are up- and down-regulated in WT versus *SETD2-*KO HKC cell lines. WT/KO protein ratios are ranked and evaluated for Gene Set Enrichment Analysis (GSEA) using the Gene Ontology datasets. Gene sets with an FDR q-value <0.25 were considered to be significant (-log_10_ = 0.60). **b)** Displayed are the ten most significant gene sets that are increased and decreased in *SETD2-*KO cells relative to wild type cells. **c)** The Gene Ontology signature with the lowest FDR q-value is GO_Translational_Initiation, which demonstrates increased levels of several proteins involved in protein translation in the *SETD2-*KO cell lines.

### TCGA ccRCC cohorts have similarly dysregulated protein translation networks

To determine if protein translation was likewise altered in *SETD2*-mutated tumors from patients, we analyzed data from the ccRCC cohort of the Cancer Genome Atlas (TCGA). As the SET domain of *SETD2* is responsible for modulating expression of EEF1AKMT2/3, the mutant group was defined as patients with a *SETD2* mutation predicted to abolish methyltransferase activity, either through a truncating mutation prior to the SET domain or a mutation within the domain (termed Mut.SET.domain) (Fig. 6a). The comparison groups were patients with wildtype SETD2 or the remaining SETD2 mutants not previously included. Mut.SET.domain patients demonstrated shorter progression free survival (PFS) and a trend towards decreased overall survival (OS) (Fig. 6b) as well as decreased expression of EEF1AKMT2 (p=0.03) (Fig. 6c). Low expression of EEF1AKMT2 is associated with poor survival in these patients (Supplemental Fig. 3)^28^. Evaluation of the TCGA gene expression data demonstrates increased expression of genes associated with protein translation in the Mut.SET.domain tumors (Fig. 6d-e), just as the SETD2-KO cell lines demonstrated increased levels of proteins (Fig. 5b, c) and genes (Supplemental Fig. 4 and Supplemental Table 3) associated with protein translation. Overall, data from both human cell lines and human tumors suggest that loss of methyltransferase activity of SETD2 is associated with increased expression of genes and proteins associated with protein translation.

**Figure 6.**
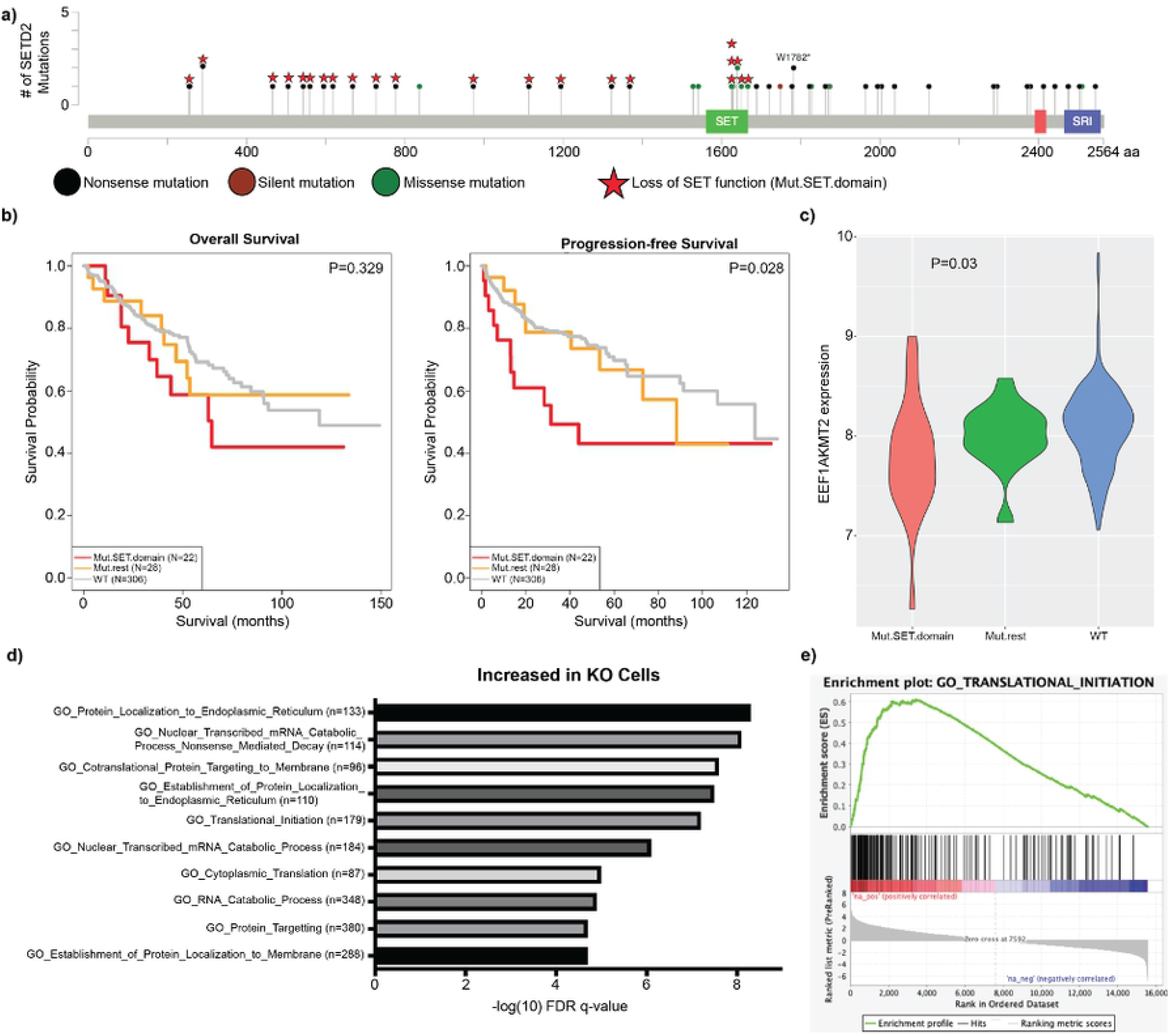
SET domain deficient ccRCC patient tumors demonstrate decreased EEF1AKMT2 expression and upregulated translation gene sets. **a)** Mutated SET domain (Mut.SET.domain) was defined as patients with either mutated SET domain or truncating mutations prior to the domain (n = 24). The remainder of patients with SETD2 mutations were classified as Mut.rest (n = 28). Patients with wildtype *SETD2* gene were labeled WT. **b)** Mut.SET.domain had significantly worse PFS than WT SETD2 patients, with a trend towards worse OS (2 patients in Mut.SET.domain were removed due to missing survival data). **c)** EEF1AKMT2 expression is decreased in tumors with SETD2 SET domain mutations (p = 0.03). **d)** Displayed are the most significantly upregulated Gene Ontology signatures in Mut.SET.domain ccRCC. **e)** Similar to SETD2-KO cell line data, protein translation signatures are upregulated in SET domain deficient ccRCC, including GO_Translational_Initiation.

## Discussion

We utilized the immunoprecipitation and purification of lysine-methylated peptides coupled with SILAC-based mass spectrometry as a method to explore how loss of *SETD2* in human kidney cells perturbs the tumor proteome. We coupled this quantitative, PTM-focused approach with systems-wide analyses of gene and total protein expression to integrate changes in the methylproteome with global cellular function. Finally, we translated our discoveries by incorporating data from human ccRCC tumors into our final analysis. The resultant data implicates a previously undescribed role of *SETD2* in the regulation of protein translation.

We show that the loss of SETD2 results in the robust decrease of trimethylation of eEF1A1 at K165 and monomethylation of K318 in multiple cell types, including a ccRCC-derived line. While SETD2 is unlikely to directly methylate eEF1A1 as it lacks the SETD2 methylation consensus sequence, our data suggest an indirect mechanism whereby SETD2 influences eEF1A1 methylation. Presumably, this decrease occurs through the decreased transcriptional expression of EEF1AKMT3 and EEF1AKMT2. While we have not functionally investigated the consequences of eEF1A1 methylation, pathway enrichment analysis reveals the upregulation of translation-associated proteins and genes with the loss of SETD2 in these cell lines. Interestingly, we similarly see an increase in expression of protein translation-associated transcripts with the loss of SETD2 catalytic activity in TCGA ccRCC cohorts, along with a decrease in EEF1AKMT2 expression. Collectively, our results indicate a possible role for SETD2 in the regulation of protein translation that warrants further investigation.

The methodology applied readily accounts for the inability to consistently measure the loss of lysine methylation of canonical SETD2 targets, such as histone H3 lysine 36 and alpha tubulin lysine 40. Lysine methylation leading to poor trypsin efficiency as well as the distribution of trypsin cleavage sites can produce peptide fragments that are suboptimal for LC-MS/MS identification and quantification. These features of the assay must also be taken into account in considering other methylation targets which may have been overlooked in this analysis.

Recent research has identified the methyltransferases and methylation sites for human eEF1A1. Currently, there are six known methylated lysines on human eEF1A1: the N-terminus, K36, K55, K79, K165, and K318^19^. The N-terminus and K55 are methylated by a single methyltransferase, METTL13, while the remainder are methylated by EEF1AKMT1-4^19,29^. Though one of the most methylated eukaryotic proteins, the consequences of eEF1A1 methylation are still poorly understood^19^. Studies have shown a range of possible effects to translation, including altering general translation rate, the specific transcripts translated, and even codon-specific rates^17,26,30^. Of the two eEF1A1 methylation states that we investigated, K165 methylation is best described. Loss of this methylation results in global reprogramming of peptide synthesis (i.e., translatome), with upregulation in translation of some mRNA molecules and down-regulation of others. Interestingly, pathway analysis of EEF1AKMT3-null cells reveals increased translation of mRNA molecules associated with protein translation, similar to what we observed in SETD2 SET domain-deficient cells^26^. This provides further evidence that SETD2 may regulate protein translation through the indirect modulation of eEF1A1 methylation.

How the various eEF1A1 lysine methyl transferases influence tumor progression is still unclear. We provide data that loss of SETD2, a tumor suppressor whose loss promotes malignant progression in ccRCC and other cancers, results in decreased expression of the methyltransferases EEF1AKTM2/3. In addition, decreased expression of EEF1AKMT2 correlates with poor survival in ccRCC. However, eEF1A1 methylation has been shown to serve as a tumor permissive PTM – albeit at a different site. Liu et al. found that Kras-driven cancers were reliant on the methylation of eEF1A1 by METTL13 at K55^17^. Loss of this methyltransferase led to decreased proliferation and viability of these malignancies due to decreased peptide synthesis. This dependence appears to be malignancy specific, as METTL13 in bladder cancer opposingly acts as a tumor suppressor^31^. Overall, further work is required to understand how these eEF1A1 lysine methylation sites coordinate in the regulation of protein translation and influence tumor progression within the context of different tumor types.

Given that eEF1A1 has noncanonical roles outside of the translatome, such as emerging roles in apoptosis and cytoskeletal regulation, decreased eEF1A1 methylation in the context of SETD2 loss could have other unrelated tumor modulating properties^32^. In fact, the loss of EEF1AKMT2, though remaining to be described in humans, affects viral replication in plants^33^. Post translational modification of eEF1A1 is a highly dynamic process that is only beginning to be uncovered^26^. Much like the histone code that affects gene transcription, protein translation may be regulated by a “translation code” composed of PTM “readers”, “writers” and “erasers” that remain to be discovered. Given the dynamics of malignancy and protein translation, combined with emerging data on the role of eEF1A methylation in cancer, this translation code warrants further research.

## Materials and Methods

### Cell Culture

The HKC cell line was obtained from Dr. Lorraine Racusen (Johns Hopkins University, Baltimore, MD)^34^. 786-0 and 293T cell lines were obtained from the American Type Culture Collection (ATCC). *SETD2*-null cell lines were previously generated using transcription activator-like effector nucleases (TALEN) targeted to exon 3 of *SETD2*, as described^20,23^. Truncated SETD2, a SET-domain mutant (R1625C), and an SRI-domain mutant (R2510H) were introduced in SETD2-deficient cells as previously described^23^. 786-0 cells were cultured in RPMI media (Gibco, catalogue no. 11875093), while HKC and 293T cells were cultured in DMEM media (Gibco, catalogue no. 11965092). All media contained 10% FBS and 1% penicillin/streptomycin. All cell lines were tested for mycoplasma contamination before use and at regular intervals during experimentation (ABM, catalogue no. G238).

### Immunoblot Analysis

Each adherent cell line was washed with 1X PBS. Cells were lysed in RIPA buffer (Sigma, catalogue no. R0278), followed with a 30-minute (min) incubation on ice. Whole cell lysate was collected after centrifugation at 20,000 g for 20 min. For each sample, 20 μg of protein was run on a 4-20% SDS-PAGE gel and transferred onto polyvinylidene difluoride (PVDF) membranes. After blocking with 5% bovine serum albumin in TBST (50 mmol/L Tris pH 7.5, 150 mmol/L NaCl, and 0.1% Tween 20), membranes were probed overnight at 4°C with anti-SETD2 (Sigma, catalogue no. HPA042451) and anti-beta-actin (Cell Signaling Technology, catalogue no. 4970) antibodies. After washing with TBST, membranes were probed with either goat anti-mouse IgG-HRP (Promega, catalogue no. W4021) or goat anti-rabbit IgG-HRP (catalogue no. W4011). ImageJ software was used for blot intensity quantification.

### Histone Extraction

Histones were acid extracted overnight as previously described (www.Abcam.com), followed by acid neutralization with Tris-HCl, pH. 8.0. Histone post-translational modifications were then analyzed using 2.5 μg of extract and anti-H3K36me3 (Cell Signaling Technology, catalogue no. 4909) and anti-pan H3 (Abcam, catalogue no. ab1791) antibodies through immunoblot analysis.

### Detection of global lysine methylation through mass spectrometry

Cells were SILAC labeled per established protocols^35,36^. Briefly, cells were cultured with either “light” media containing lysine and arginine with normal isotopic distribution or “heavy” media containing 13C/15N-isotope labeled lysine and arginine. Cells were passaged 10 times and SILAC labelling efficiency checked via mass spectrometry prior to further analysis. Whole protein lysates were analyzed for total protein expression and lysine-methylated peptides were enriched using reagents from Cell Signaling Technology (catalogue no. 14809). In short, lysate from SILAC-labelled isogenic SETD2 WT and KO HKC whole cell lysates were trypsin-digested, then immunoprecipated with anti-pan-methyl lysine antibody beads. Peptide eluate was then identified and quantified via liquid chromatography column and tandem mass spectrometer (LC-MS/MS) as described previously^37^. For whole proteome analysis, aliquots of the mixed SILAC-labeled lysates containing 20 μg of protein were acetone precipitated, reduced, alkylated, and trypsin-digested. LC-MS/MS analysis of the peptides was performed using a Q Exactive mass spectrometer (Thermo Scientific). The peptides were loaded onto a self-packed biphasic C18/SCX MudPIT column using a Helium-pressurized cell (pressure bomb). LC-MS/MS was performed with an 11-step ammonium acetate salt pulse gradient. Peptides were eluted from the analytical column after each salt step with a 90-min reverse gradient (2-50% acetonitrile, 0.1% formic), followed by a 10-min equilibration at 2 % B, for the first 10 salt pulses. For the final salt step, a gradient consisting of 2-98% acetonitrile was used. Data were collected using a data-dependent method. Please see Supplemental Methods for further detail.

### Mass spectrometry and pathway enrichment analyses

For both total protein and lysine-methylated peptide proteomics data analysis, the raw files were processed with the MaxQuant v1.5.8.3^38^ and searched with Andromeda search engine against the human UniProt database v20120610 (20245 entries) and integrated with FASTA sequences of 247 contamination proteins. The precursor ion mass deviations of first search and main search were set as 20 ppm and 10 ppm, while fragment mass deviation of 20 ppm was used. The minimum peptide length was set to 7 amino acids and strict specificity for trypsin cleavage was required, allowing up to two missed cleavage sites. Multiplicity of 2 was used, selecting arginine (+10 Da) and lysine (+8 Da) as heavy labels. Re-quantification and matching between runs were selected. Protein was identified with a minimum of one unique peptide. The false discovery rates at the protein, peptide and site level were set to 2%, 1% and 2%. Oxidation (M) was set as variable modification and Carbamidomethyl (C) was set as static modification. For the lysine-methylated peptide dataset, Methyl (K), Dimethyl (K) and Trimethyl (K) were also set as variable modifications. The chromatograph of identified methylated peptides were extracted using ProteomicsTools and manually validated based on light/heavy ion chromatograph correlation, distance between theoretical and observed peptide mass profile, and light/heavy isotopic profile cosine similarity^39^. For total proteomics dataset, quantified proteins were ranked by mean of log ratio cross samples and gene set enrichment analysis was performed using GSEA package^40^. Then, the proteins quantified in only one sample or the proteins with inconsistent relative quantification change direction (knockout vs wild type) were discarded. Genome Ontology and KEGG pathway over-representation analysis was performed on differentially expressed proteins with fold change larger than 1.5 in at least one sample using the WebGestaltR package.

For RNA sequencing data, reads were aligned to the human b37 genome using STAR v2.5.3a^41^. Ensembl v75 gene annotations were provided to STAR to improve the accuracy of mapping. Quality control on raw reads was performed using FastQC (www.bioinformatics.babraham.ac.uk/projects/fastqc). FeatureCounts v1.15.2^42^ was used to count the number of mapped reads to each gene. Significantly differential expressed genes with FDR-adjusted p-value < 0.05 and absolute fold change > 2.0 were detected by DESeq2 (v1.18.1)^43^. Heatmap3^44^ was used for cluster analysis and visualization. Genome Ontology and KEGG pathway over-representation analysis was performed on differentially expressed genes using the WebGestaltR package^45^. Gene set enrichment analysis was performed using GSEA package^40^.

### Size enrichment of eEF1A1 proteins

SILAC-labeled protein lysates from SETD2-WT and -KO cells were mixed 1:1, and 50 μg of the mixed lysates (25 μg of WT lysate and 25 μg KO lysate) were loaded onto a NuPAGE 10% Bis-Tris gel. The gel was stained with Coomassie stain, and the region corresponding to approximately 48 – 52 kDa was excised and diced for in-gel digestion. Proteins were reduced, carbamidomethylated, destained, and trypsin digested. Peptides were extracted by gel dehydration, dried by speed vac centrifugation, and reconstituted in 0.1% formic acid and analyzed by LC-coupled tandem mass spectrometry (LC-MS/MS) via methods similar to those previously described. Peptides mass analysis was performed using a data-dependent method, with an inclusion list of specific m/z values corresponding to various forms of the eEF1A1 peptide MDSTEPPYSWKR. The inclusion list included methylated forms of the peptide in both light and heavy labeled conditions. Comparison of the heavy and light forms of eEF1A1 MDSTEPPYSQK(me3)R (eEF1A1_K318) peptide were conducted in WT and KO cells in two replicates independent of lysine methyl-peptide enrichment. See Supplemental Methods for further detail.

### Synthetic heavy-labeled peptide-aided peptide quantification

A synthetic peptide labeled with heavy arginine (13C, 15N) was formulated for the MDSTEPPYSQK(me3)R (eEF1A1_K318) peptide (HeavyPeptide AQUA custom synthesis service, Life Technologies). Synthetic peptides were spiked into the WT and *SETD2-*KO lysates from the HKC and 786-O cells, allowing for relative quantification of the endogenous methylated peptide in non-SILAC-labeled cells. First, HKC and 786-O lysates (50 μg) were separated on gel as described previously, and the eEF1A1 gel regions were excised and in-gel digested with trypsin. WT and KO lysates were prepared in triplicate for LC-MS/MS analysis of eEF1A1. Following in-gel digestion, the peptides were reconstituted in 20 μL of 0.1% formic acid. Aliquots (8 μL) of each digest from the triplicate WT and KO lysates were then spiked with the synthetic peptide to make a solution of 12 μL containing 50 fmol/μL of the synthetic peptide. For LC-MS/MS, 2.5 μL of the spiked in-gel digests were analyzed on a Q Exactive Plus mass spectrometer. The method consisted of both data-dependent and targeted PRM scan events. Following data-dependent MS2 scan events, the method included targeted PRM scans of m/z values corresponding to the light and heavy eEF1A1 peptide MDSTEPPYSWK(me3)R. Targeted m/z values included oxidized and unoxidized forms of the peptide in both light and heavy SILAC states. PRM data were imported into Skyline^46^, product ions were evaluated, and integrated areas were calculated in Skyline for y-type ions, y6 - y11, for each peptide precursor. Areas were summed for the light precursors and heavy precursors separately, and then ratios of the summed areas for WT and KO samples were calculated and used to determine the difference in the relative amount of the peptide MDSTEPPYSWK(me3)R peptide. See Supplemental Methods for more detail.

### RT-qPCR

Cells were harvested and RNA was extracted using the RNeasy Mini Kit (Qiagen, catalogue no. 74106). For each sample, a total of 500 ng of RNA was reverse transcribed into cDNA using oligo(dT) from the SuperScript IV First-Strand Synthesis System (Thermo Fisher, catalogue no. 18091050). After completion of cDNA synthesis, each reaction was diluted 1:10. qPCR was performed in triplicate, using 10 μL of SYBR Green PCR Master Mix (Thermo Fisher, catalogue no. 4309155), 4 uL of diluted cDNA, 1 μL of 10 uM primers, and 5 μL of nuclease-free water.

PCR plates were run on a CFX96 Touch Real-Time PCR Detection System (BioRad), with a program of 10 min at 95°C and 40 cycles of 95°C for 10 seconds and 62°C for 30 seconds. Amplification purity was tested by melt curve. Real-time PCR data was analyzed using the comparative C_T_ method^47^.

### Patient cohort analyses

TCGA kidney renal cell cancer datasets were downloaded from cbioportal.org. SET domain mutations were defined as non-synonymous mutations in the SET domain of *SETD2*, or truncating mutations in the 5’ region of the SET domain. Differentially expressed genes associated with SET domain mutations vs. WT samples were identified using t-test, and gene set enrichment analysis was performed using GSEA using MSigDB C5 genesets^40^. Analyses were performed using R version 3.6.0.

---

**Supplemental Figure 1. Immunoblots validating loss of SETD2 protein and catalytic activity. a)** *SETD2* was knocked out of HKC cells using TALEN. Successful knock out was confirmed through immunoblot analysis. Functional loss of SETD2 was further validated by H3K36me3 immunoblot. Both SETD2 and H3K36me3 were decreased relative to loading control in *SETD2*-KO cell line (relative quantifications shown). **b)** Validation of rescue with the truncated SETD2 variants by SETD2 and H3K36me3 immunoblot (relative quantifications shown). **c)** Uncut films of SETD2-WT, KO, tSETD2, SET-mt, and SRI-mt cell lines. The rescue vectors included a flag tag, showing successful construct expression.

**Supplemental Figure 2. Distribution of heavy isotope-labeled (H) and unlabeled (L) peptides in each replicate (standard and normalized)**. Normalized ratios were used for all subsequent analyses.

**Supplemental Table 1. Lysine-methylated peptides in WT and *SETD2-*KO human kidney cell lines**. “Position” corresponds to location of methylated lysine residue within the full-length protein. “MeK” indicates whether the lysine residue was mono-, di-, or tri-methylated. When more than one lysine was methylated within a peptide, “Methyl K Probabilities” indicates the calculated relative probability that the indicated lysine residue was modified. A-D indicates the four biological mass spectrometry replicates and the ratio of the wild type (WT) and *SETD2-* knock out (KO) ratios (i.e., WT/KO ratio). If the lysine predicted to be modified corresponds to a canonical *SETD2* methylation motif (e.g., KxP or KxxG), it was indicated with “x”. “K motif” corresponds to the modified lysine and the four amino acids that follow (N-terminal to C-terminal).

**Supplemental Table 2. Differentially expressed proteins in WT and *SETD2-*KO human kidney cell lines**. Proteins quantified in only one sample or proteins with inconsistent relative quantification change direction (knockout vs wild type) were discarded. Differentially expressed proteins with fold change larger than 1.5 in at least one sample were included in pathway analysis.

**Supplemental Figure 3. Low expression of EEF1AKMT2 in ccRCC correlates with poorer survival (P=0**.**01)**. Figure obtained from UALCAN (Chandrashekar et al., 2017).

**Supplemental Figure 4. Total Transcriptome Analysis Demonstrates Changes in Protein Translation in *SETD2-*KO Cells. a)** Individual genes are up- and down-regulated in WT versus *SETD2-*KO HKC cell lines. WT/KO protein ratios are ranked and evaluated for Gene Set Enrichment Analysis (GSEA) using the Gene Ontology datasets. Gene sets with an FDR q-value <0.25 were considered to be significant (-log_10_ = 0.60). Displayed are the ten most significant gene sets that are **b)** increased and **c)** decreased in *SETD2-*KO cells relative to wild type cells. **d)** The Gene Ontology signature is GO_Translational_Initiation was highly significant and up-regulated in *SETD2-*KO cells, just as the it was in the total protein dataset (Fig. 5b). Increased levels of several genes involved in protein translation were observed in the *SETD2-*KO cell lines. **e)** The Gene Ontology signature with the lowest FDR q-value that is decreased in *SETD2-*KO cells is GO_SWI_SNF_Superfamily_Type_Complex though it does not reach the significance threshold.

**Supplemental Table 3. Differentially expressed genes in WT and *SETD2-*KO human kidney cell lines**. Differential gene expression is expressed as log2 fold change of WT/KO cell lines (three replicates for each genotype).

